# β2-Adrenergic Biased Agonist Nebivolol Inhibits the Development of Th17 and the Response of Memory Th17 Cells in an NF-κB-Dependent Manner

**DOI:** 10.1101/2024.09.08.611829

**Authors:** Mehri Hajiaghayi, Fatemeh Gholizadeh, Eric Han, Samuel R. Little, Niloufar Rahbari, Isabella Ardila, Carolina Lopez Naranjo, Kasra Tehranimeh, Steve C.C. Shih, Peter J. Darlington

## Abstract

**Introduction:** Adrenergic receptors regulate metabolic, cardiovascular and immunological functions in response to the sympathetic nervous system. The effect of β2-adrenergic receptor (AR) as a high-expression receptor on different subpopulations of T cells is complex and varies depending on the type of ligand and context. While traditional β2-AR agonists generally suppress T cells, they potentially enhance IL-17A production by Th17 cells. The effects of pharmacological drugs that count as biased agonists of AR like nebivolol are not completely understood. We investigated the impact of nebivolol on human memory CD4+ T (Th1, Th2, Th17) cells and polarized naïve Th17 cells highlighting its potential for IL-17A suppression via a non-canonical β2AR cell-signaling pathway.

**Methods:** The effects of nebivolol were tested on healthy human peripheral blood mononuclear cells, purified memory Th cells, and polarized naïve Th17 cells activated with antiCD3/antiCD28/antiCD2 ImmunoCult reagent. IFN-γ, IL-4, and IL-17A which are primarily derived from Th1, Th2, and Th17 cells respectively, were quantified by ELISA and flow cytometry. IL-10 was measured by ELISA. Gene expression of RORC, ADRB1, ADRB2, and ADRB3 was evaluated by qPCR. The ADRB2 gene was knocked out in memory Th cells using CRISPR/Cas9. Protein expression of phosphorylated-serine133-CREB and phosphorylated-NF-κB p65 was assessed by Western blot. Proliferation was assessed by fluorescent dye loading and flow cytometry.

**Results:** Nebivolol treatment decreased IL-17A and IFN-γ secretion by activated-memory Th cells and elevated IL-4 levels. Nebivolol reduced the proportion of IL-17A+ Th cells and downregulated RORC expression. Unlike the β2-AR agonist terbutaline, nebivolol inhibited the shift of naïve CD4+ T cells towards the Th17 phenotype. IL-10 and proliferation index remained unchanged. Nebivolol-treated β2-knockout memory Th cells showed significant inhibition of β2AR-mediated signaling, evidenced by the absence of IL-17A suppression compared to controls. Phosphorylation of the NF-κB p65 subunit was inhibited by nebivolol, but CREB phosphorylation was not changed, suggesting a selective transcriptional control.

**Conclusions:** The findings demonstrate that nebivolol acts through a β2-AR-mediated signaling pathway, as a distinctive anti-inflammatory agent capable of selectively shifting Th17 cells and suppressing phosphorylation of NF-κB. This highlights nebivolol’s potential for therapeutic interventions in chronic autoimmune conditions with elevated IL-17A levels.

## 1. Introduction

The sympathetic nervous system (SNS) is best known for its role in the fight or flight response where the catecholamines epinephrine and norepinephrine (NE) modulate metabolic, cardiovascular and immune systems (1-3). Complex interactions regulate adaptive immune functions through neuro-endocrine mechanisms (4). In the spleen and lymph nodes, sympathetic nerve fibres run alongside the location where T cells and dendritic cells (DC) reside, indicating the potential neural modulation of T cell functions (5-7). NE acts on T cells through adrenergic receptors (AR). Several of the AR family members are classical G protein-coupled receptors (GPCRs). Upon stimulation, β2-AR initiates a canonical pathway involving Gs proteins, leading to adenylyl cyclase (AC) activation, increase in Camp production, and phosphorylation of cAMP response element-binding protein (CREB) on Ser133 by PKA (8, 9). These processes modulate T helper cell activation, proliferation and production of certain cytokines including IL-2 and IFN-γ (10, 11). Receptor desensitization and signal termination occur through phosphorylation by regulatory kinases, such as GPCR kinases (GRKs), followed by β-arrestin-1 activation at Ser412 (8, 12). β2-ARs also engage in non-canonical, G protein-independent signaling via β-arrestin2 which leads to sustained ERK1/2 activation (13). These pathways could regulate immune responses during chronic sympathetic tone by distinct downstream effects, including modulation of gene expression and inhibition of NF-κB activation (14). NF-κB is widely considered to be a pro-inflammatory transcription factor that drives the expression of IFN-γ and IL-17A in T cells, its activity can be measured by assessing the phosphorylation of NF-κB p65 subunit at position Ser529 (15, 16).

Accumulating evidence supports a role for the β2-AR subtype in regulating T cells. Human CD4^+^ T (T helper) cells and CD8^+^ T (T cytotoxic) cells exhibit a predominant expression of β2-AR (17). Memory T cells, crucial for faster and more effective immune responses upon antigen re-exposure, exhibit increased β2AR expression compared to naïve T cells, suggesting heightened responsiveness to immune and gene expression changes induced by the SNS (18, 19). Th1 cells produce IFN-γ and express the transcription factor *TBX21*, Th17 cells produce IL-17A, among other cytokines, and express the transcription factor ROR*γ (RORC)*, and Th2 cells express IL-4, IL-5 and express the transcription factor *GATA3 (*20). Activation of naïve CD4^+^ T cells in the presence of NE and IL-12 *in vivo* induced more IFN-γ and polarized the response towards Th1 (21). β2-ARs are also expressed on dendritic cells, and NE treatment reduced IL-12 secretion, altering the IL-12/IL-23 balance. As a result, stimulated DCs promoted T helper cells to produce high IL-17A and lower IFN-γ (22). Consistent findings were observed by stimulating β2-AR with salbutamol in murine bone marrow-derived DC, which secreted IL-6, IL-23, and upregulated the Th17 response (23). Similarly, salbutamol-treated DCs reduced the secretion of Th2 cytokine IL-4 (24). Our reports demonstrated that a specific β2-agonist drug (terbutaline) significantly increased IL-17A levels in a PKA-dependent manner in human memory Th cells (25, 26). Therefore, agonists of the β2-ARs may shift the balance of Th1, Th2 and Th17 cells by promoting a Th1 and/or Th17 bias in memory Th cells. Furthermore, it may alter the polarization of naïve T cells as they shift to Th17 effector cells.

Emerging studies on β2-biased agonist drugs showed anti-inflammatory properties (27) by increasing the serum levels of IL-10, which is mainly produced by regulatory T cells (Treg) (28) and inhibited IL-2 production in activated human T cells (29). Biased agonists are ligands that preferentially activate one pathway over another, leading to distinct signaling outcomes compared to traditional agonists. Nebivolol is a third-generation β-blocker that is highly selective for β1AR (30). Moreover, it can induce nitric oxide through the β3AR (31). Nebivolol has a prominent pharmacological mechanism involving a β-arrestin biased agonist at the β2AR (32) and it is FDA-approved to treat hypertension and heart failure (33, 34), but its impact on the immune system is not completely understood. We tested nebivolol on human memory Th1, Th2, Th17, and IL-10, and differentiated naïve Th cells into Th17 cells, to determine if it shifted the immune response toward an anti-inflammatory response through β2AR cell-signaling pathways.

## 2. Materials and Methods

### 2.1. Isolation of human Peripheral Blood Mononuclear Cells (PBMCs)

Blood was collected from healthy participants after an informed, signed consent had been provided. Concordia University Human Research Ethics Committee, which follows the Declaration of Helsinki guidelines, approved the study (certificate 30009292). Health was determined by self-reporting during a semi-structured interview. Exclusion criteria included those under 18, who have specific medical conditions, or take ineligible medications. If a participant took recreational drugs or was vaccinated in the past 2 weeks their blood draw was rescheduled. 1 X PBS was used to dilute heparinized peripheral blood at a 1:1 V/V In 50-ml conical tubes, 30 mL of diluted blood was layered over 15 mL of lymphocyte separation solution (Wisent Bioproducts, CA). The sample was centrifuged at 624g for 30 min at room temperature (RT). In a separate 50-ml tube, mononuclear cells were collected and centrifuged at 433g for 15 min with 25 ml of PBS. The supernatant was discarded, and the pellet was resuspended in 25 ml of PBS and centrifuged at 400 g for 12 minutes again. In order to isolate the T cells from the cell pellet, PBS containing 2% heat-inactivated fetal bovine serum (FBS) was used as a resuspension solution.

### 2.2. Isolation, and purification check of CD4+ naïve and memory T cells

Naïve (CD3^+^ CD4^+^ CD45RA^+^ CD45RO^-^) and memory (CD3^+^ CD4^+^ CD45RA^−^ CD45RO^+^) T cells were isolated from PBMCs in the recommended medium (PBS containing 2% FBS and 1 mM EDTA) by using EasySep® Human Naïve CD4^+^ T Cell Isolation Kit II and human memory CD4^+^ T cell enrichment kit respectively (StemCell Technologies) according to the manufacturer’s instructions. Purity was analyzed by flow cytometry (FACS Verse, BD Biosciences) using CD3-Peridinin-chlorophyll-Cyanine5.5 (BD Biosciences, clone OKT3), CD4-allophycocyanin (BD Biosciences, clone RPA-t4), anti-CD45RA-Fluorescein isothiocyanate (BD Biosciences, clone HI100), and anti-CD45RO-phycoerythrin (BD Biosciences, clone UCHL1) mAbs. Purity was found to be >90% (Supplementary figures).

### 2.3. Stimulation, culture, and drug treatment of memory Th cells

Purified memory Th cells were suspended in RPMI 1640 medium (Wisent Bioproducts) containing 10% heat-inactivated FBS, 1% penicillin and streptomycin and 1% Glutaplus (Wisent Bioproducts) at the final concentration of 0.3×10^6^ cells/ml and stimulated with ImmunoCult™ Human CD2/CD3/CD28 T Cell Activator (Stemcell Technologies). In the next step, nebivolol hydrochloride (Sigma Aldrich, CA) dissolved in glycerol, was added to the purified memory Th cells and incubated the plate for 5 days in 96-well U-bottom plates at a final concentration of 10 μM. Since nebivolol was dissolved in glycerol, we included vehicle control groups with only glycerol added at an equivalent dilution. The activated memory Th cells were also treated with another β_2_-biased agonist carvedilol (10 μM, Sigma Aldrich, CA) or combined with a selective βI□ receptor antagonist metoprolol tartrate (10 μM, Sigma Aldrich, CA) or a non-selective β adrenergic receptors antagonist bupranolol (10 μM, Cayman Chemical, CA) and H89 as a PKA inhibitor (2 μM, Sigma Aldrich, CA) 30 minutes before adding the activator.

### 2.4. In vitro Th17 cell polarization and treatment of CD4+ naïve T cells

Naïve CD4^+^ T cells were cultured at 0.3□×□10^6^ cells/well in X-VIVO™ 15 serum-free hematopoietic cell medium (Lonza, USA). Cells were expanded with ImmunoCult™ human CD2/CD3/CD28 T cell activator (Stemcell Technologies). Four types of cultures were established: (1) nonpolarizing conditions as a control, (2) Th17-polarizing conditions in the presence of human recombinant TGF-β (1 ng/ml;

Cedarlane, CA), human recombinant IL-21 (25 ng/ml; Cedarlane, CA), human recombinant IL-23 (25 ng/ml; Cedarlane, CA), anti-IL-4 (10 μg/ml; Abcepta, CA), and anti-IFN-γ (10 μg/ml; Abcepta, CA), (3) Th17-polarizing conditions as above plus 10 μM of nebivolol, and (4) Th17-polarizing conditions with 10 μM of terbutaline. After 7 days of culture, the following procedures were performed.

### 2.5. Cytokine analysis using ELISA

The culture supernatants of either PBMC cells, memory Th cells, or polarized Th17 cells were collected after incubation by centrifugation of the 96 well culture plate at 100g for 6 minutes to remove any debris. The supernatant was analyzed for IL-4, IFN-γ, IL-10 (BD Biosciences); and IL-17A (ThermoFisher Scientific) with ELISA kits following the manufacturer’s instructions. A standard curve was generated for each assay, with the limits of detection for IL-17A, IL-4, and IL-10 = 500 pg/ml, and IFN-γ= 300 pg/ml. Supernatants were diluted accordingly with assay diluent. Two technical ELISA replicates of the standard curve and each experimental group were done. Data was acquired on a spectrophotometer at 450nm with a 570nm correction factor (Biotek Agilent, Fisher Scientific).

### 2.6. Flow cytometry

Intracellular IL-17A and IFN-γ were measured using established intracellular cytokine staining (ICS) procedures (35). In brief, after cell culture was complete, the cells were incubated with 0.5μg/ml of ionomycin, 10 μg/ml of brefeldin A and 0.05 μg/ml of phorbol 12-myristate 13-acetate (Sigma Aldrich Millipore, Oakville, Canada) for four-hours at 37 □C at 5% CO_2_. The cells were fixed and permeabilized with a kit following the manufacturer’s instructions (BD Biosciences, CA). Cells were incubated with saturating concentrations of fluorochrome-labeled Abs. The antibodies were CD4-allophycocyanin (BD Biosciences, clone RPA-t4), IL-17A-phycoerythrin (R & D Systems, clone 41802), INF-γ-Brilliant Violet 421 (BD Bioscience, clone B27), IL-4-phycoerythrin-Cyanine7 (BD Biosciences, clone 8D4-8). Samples were incubated for 45 min, on ice in the dark. Two washes (493 x g, 5 minutes) with a staining buffer (1% FBS in 1X phosphate buffered saline) were done and the samples were resuspended in a 500 μl staining buffer in microcentrifuge tubes. Each set of runs consisted of three technical replicates of each experimental group. At least 50,000 events were obtained on the flow cytometer (FACS Verse, BD Bioscience), and analyzed with FlowJo software (BD Biosciences).

### 2.7. Cell-proliferation and viability assessment by using CFDASE and 7AAD staining

According to a published protocol (26), the tracking dye 5(6)-Carboxyfluorescein diacetate N-succinimidyl ester (CFDASE) was used to measure proliferation. Memory Th cells were labelled with CFDASE, incubated in the specified activation conditions for five days, and then stained for a surface marker with CD4-APC. After 30 min, cells were washed and 5 μL of 7AAD-Peridinin-Chlorophyll-Protein reagent (BD Biosciences) was added to the cell suspension and incubated in the dark for 15 minutes at room temperature. In each group, at least 50,000 cells in three replicates were analyzed with a flow cytometer (BD Bioscience, CA, FACS Verse). The division index, proliferation index, and viability were determined in CD4-gated cells using the cell proliferation tool (BD Biosciences).

### 2.8. mRNA extraction and quantitative real-time PCR

The qPCR analysis was done according to published protocols (26). Briefly, memory Th cells or polarized naïve Th17 cells were cultured with activating and drug conditions. At least 2 x 10^6^ cells were collected from each experimental group, and total RNA was extracted using the PureLink™ RNA Mini Kit (BD Biosciences). The RNA concentration and purity were measured using a spectrophotometer (NanoDrop™ 2000c, ThermoScientific). The RNA samples were then converted to cDNA using iScript™ Reverse Transcription Supermix for RT-qPCR (Bio-Rad Laboratories, USA). At least two technical replicates of each independent sample were used for real-time quantitative Polymerase Chain Reaction (qPCR) using TaqMan™ gene probes from ThermoFisher Scientific to measure *ADRB1* (Hs02330048), *ADRB2* (Hs00240532), *ADRB3* (Hs_00609046_m1), *RORC* (Hs01076112) and housekeeping genes *PPIA* (Hs99999904_m1). The housekeeping gene was normalized to the average of their expression and shown as a fold increase in relative expression 2-ΔCT. Fold change is calculated as the ratio of the normalized expression level of the target gene in the treated sample to that in the activated control group which is set to 1.

### 2.9. Knocking Out *ADRB*2 Gene in Memory Th Cells Using the CRISPR/Cas9-based gene editing

Memory Th cells were activated at a concentration of 1 × 10^6^ cells/ml using Dynabeads™ Human T-Activator CD3/CD28 (Fisher Scientific Ottawa, ON, #11131D) at a 1:1 cell to bead ratio in RPMI-1640 medium supplemented with 10% FBS, 100 U/ml penicillin/streptomycin, and 100 IU/ml recombinant human IL-2 (Fisher Scientific Ottawa, ON). Cells were cultured at 37°C, 5% CO2 for two days. After incubation, activator beads were removed by gently pipetting and using a magnetic tube rack for 1–2 minutes to separate cells from the beads. The supernatant containing the cells was transferred to a fresh tube and maintained at 1 × 10^6^ cells/ml with daily addition of complete culture media.

According to published protocols (36, 37), before electroporation briefly, 100 pmol of optimized multi single-guide RNA (sgRNA) (payload sequences attached in supplementary table1) (Synthego, USA) and 50 pmol of Cas9 (Thermofisher, CA) per 1 million cells were mixed in 5 μL of Resuspension Buffer R and incubated at room temperature for 10 minutes to form the Cas9-ribonucleoprotein (RNP) complex. T cells (2 × 10^6^) were washed with PBS, resuspended in Buffer R, and mixed with Cas9/gRNA complex. Electroporation was performed using the Neon™ 100-μL tip and program #24 (1600 V, 10 ms, 3 pulses). Post-electroporation, the cells were incubated in a Recovery Buffer containing 10% FBS, 100 U/ml penicillin/streptomycin, 400 IU/ml recombinant human IL-2 at 37°C, and 5% CO2 for 72 hours. Editing efficiency was assessed using *ADRB2* mRNA expression levels via RT-qPCR in non-electroporated, non-targeting gRNA (nt-gRNA), T cell receptor alpha constant (TRAC) multi-sgRNA (Positive Control), and *ADRB2* multi-sgRNA electroporated conditions. Following that, cells were treated with nebivolol at a final concentration of 10 μM for 5 days to assess the impact of β2AR gene depletion on IL-17A production.

### 2.10. Western blot

A published protocol was followed to prepare whole-cell lysates after treatment of activated memory Th cells with nebivolol for 15 minutes (26). Protein concentrations were determined with a Bradford assay kit (BioRad). For separation by electrophoresis, 15 μg total protein in two technical replicates of each sample was loaded onto SDS-polyacrylamide gels according to standard protocols (SDS-PAGE) and then transferred to nitrocellulose. Membranes were blocked with 5% (wt/vol) non-fat milk (Anatol Spices, Montreal Canada) in Tris-buffered saline with 0.1% Tween-20 (TBST) overnight at 4°C and then again incubated overnight at 4°C with recommended dilutions of primary antibodies. Antibodies were against human Phospho-ser529 NF-κB p65 (1/500, clone A21012B Biolegend), NF-κB p65 Antibody (1/1000, clone 14G10A21, Biolegend), phospho-Ser133 CREB1 (1/700, rabbit polyclonal, Cusabio, Cedarlane), phospho-Ser412-β-arrestin1 (1/1000, clone mAb #2416 Cell Signaling Tech., Danvers, USA), β2AR (1/500, clone 4A6C9, Novusbio, Cedarlane), GAPDH (1/15000, 10B4E3, MA000071M1m, Cusabio, Cedarlane) and α□tubulin (1/1000, Santa Cruz Biotechnology Inc., USA). Subsequently, membranes were repeatedly washed with TBST and incubated for 2 h with the appropriate HRP-conjugated secondary mouse antibody (1/1500, BioRad Laboratories) or secondary goat anti□rabbit IgG HRP (1/5,000 Cusabio Cedarlane) in 5% skim milk. Immunoreactivity was detected using the ECL prime reagent (GE Healthcare), and then the chemiluminescence signal was recorded in the Image Lab (Bio-Rad Laboratories). Data were analyzed with Image Lab software (Bio-Rad Laboratories). Total α-tubulin levels were used as a loading control.

### 2.10. Statistical analysis

Statistical analyses were conducted using ANOVA with Tukey’s multiple comparison tests to assess the effects of treatment and culturing conditions on the studied parameters. All data are presented as mean ± SEM. Statistical significance was determined using p-value thresholds: *<0.05, **<0.01, ***p<0.001, and ****p<0.0001. The number of independent biological experiments and technical replicates for each experimental condition are detailed in the legend of each figure. All analyses were performed using GraphPad Prism 9 software.

## Results

### 3.1. Nebivolol promotes an anti-inflammatory shift in T helper cells, effectively suppressing Th1 and Th17 cells

To test the effects of nebivolol on Th1, Th2 and Th17 cytokines, activated human PBMCs were used in cell culture experiments. Nebivolol had a trend of decreasing IFN-γ, showed a trend of increasing IL-4, and significantly decreased IL-17A when compared to the control group without treatment and vehicle control (Figure 1A-C). In contrast, our findings indicate that carvedilol, another β2-biased agonist, did not significantly alter cytokine production (Figure 1A-C). Measurements from PBMC experiments confirmed that nebivolol at the concentration of 10 μM did not cause cellular cytotoxicity (Figure 1D). In the next step, purified memory CD3^+^CD4^+^CD45RA^-^CD45RO^+^ T cells were extracted from PBMC to a high degree of purity (Supplemental figure). Memory Th cells were activated for five days with antiCD3/antiCD28/antiCD2, with or without nebivolol treatment. Nebivolol-treated cells showed a trend of decreased IFN-γ and IL-17A production while increasing IL-4 levels compared to the activated control group without treatment. There was no significant change in IL-10 levels between the treated and untreated groups after activation (Figure 2A-D). To underscore nebivolol’s ability to shift cytokine secretion, intracellular cytokine staining was conducted on memory Th cells (Figure 3A-D). Nebivolol exhibited a trend in decreasing the proportion of IFN-γ^+^, a trend in increasing the proportion of IL-4^+^, and a significant decrease in the proportion of IL-17A^+^ memory Th cells (Figure 3E-G). Despite the changes in cytokines IFN-γ and IL-4, there were no changes in TBX21 and GATA3 as the main transcription factors of Th1 and Th2 respectively in response to nebivolol (Figure 3H, I). However, the inhibition of IL-17A was concurrent with mRNA levels since nebivolol inhibited the expression of *RORC* which is a main transcription factor of Th17 cells (Figure 3J). Together, the data indicates that nebivolol causes an anti-inflammatory shift, suggesting that it inhibits the Th17 development pathway.

**Fig 1.**
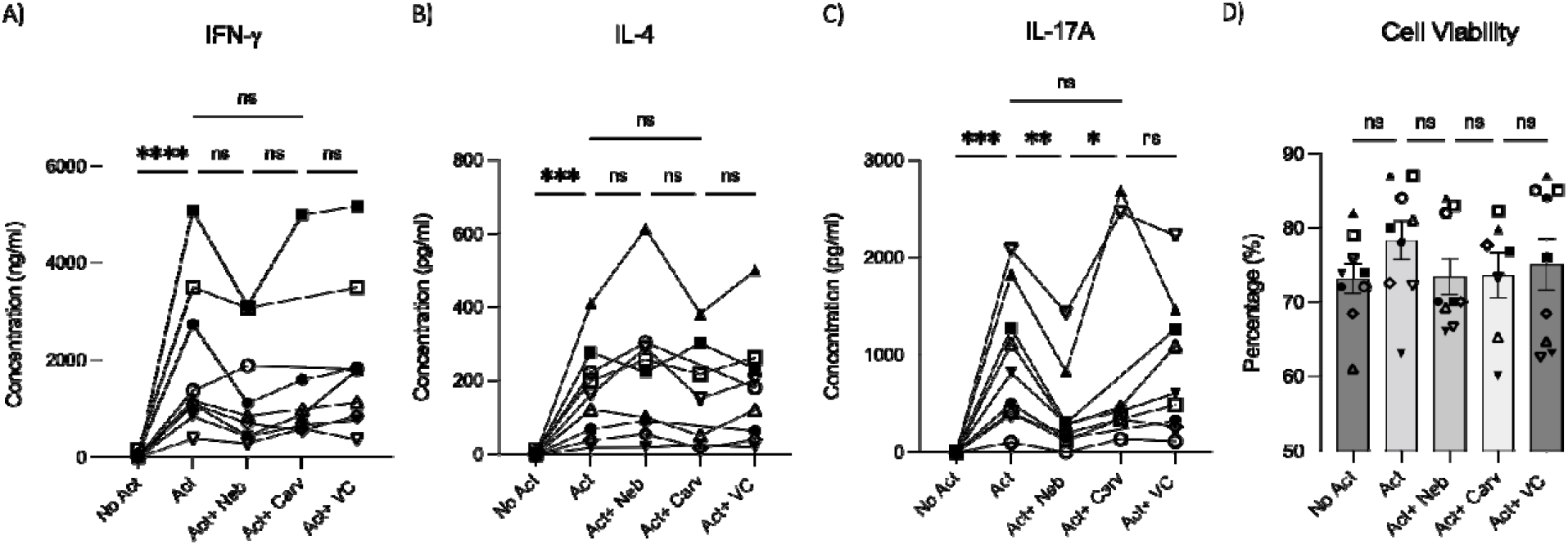
IFN-γ, IL-4 and IL-17A secretion by PBMCs treated with nebivolol or carvedilol. PBMC samples were not activated or activated with antiCD3/antiCD28/antiCD2(Act) for four days with nebivolol (Neb) or carvedilol (Carv) in the concentration of 10 μM as compared to the equivalent dilution of vehicle control (VC). The A) IFN-γ, B) IL-4, C) IL-17A levels, and D) cell viability in cell culture supernatants were measured by ELISA. The viability of the cells was measured by trypan blue. Pooled data are expressed as Mean ± SEM of seven to nine independent biological experiments. One-way ANOVA followed by Tukey’s multiple comparisons tests was performed.

**Fig 2.**
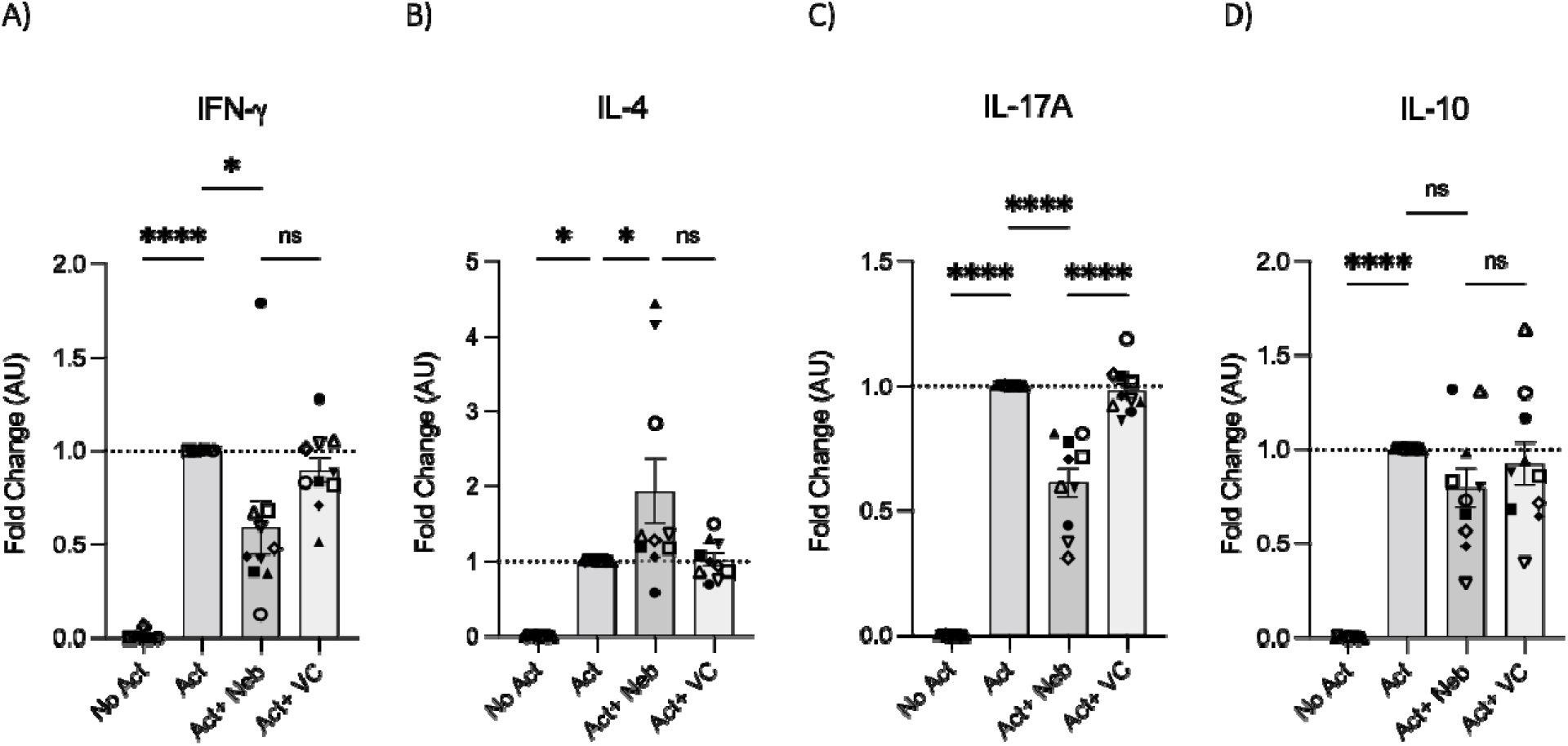
IFN-γ, IL-4, IL-17A, and IL-10 secretion by memory Th cells treated with nebivolol. Memory Th cells were not activated or activated with antiCD3/antiCD28/antiCD2 for five days and the A) IFN-γ, B) IL-4, C) IL-17A, and D) IL-10 levels in cell culture supernatants were measured by ELISA and expressed as fold change compared to the Act group, which was set to 1.0 (dotted line). Pooled data are expressed as Mean ± SEM of ten independent biological experiments. One-way ANOVA followed by Tukey’s multiple comparisons tests.

**Fig 3.**
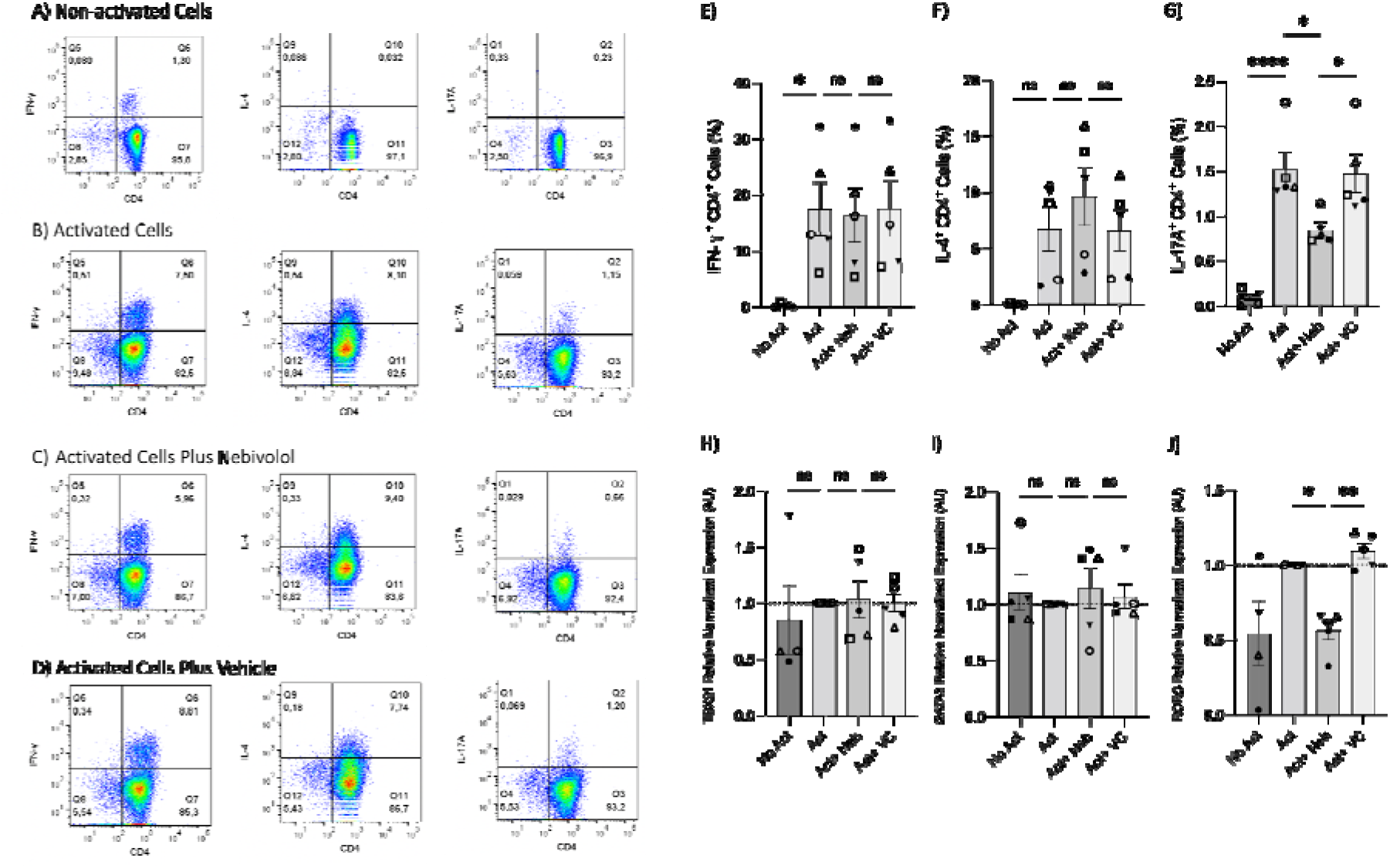
Intracellular cytokine staining and expression of transcription factor genes associated with Th1, Th2, and Th17 in memory Th cells treated with nebivolol. Memory Th cells were not activated, or activated with antiCD3/antiCD28/antiCD2 for five days and the samples were stained for intracellular cytokines with or without nebivolol treatment and analyzed by flow cytometry. Representative dot plots are shown for CD4 and each of IFN-γ, IL-4, and IL-17A antibodies on memory Th lymphocytes. A) Non-activated cells, B) Activated cells, C) Activated cells plus nebivolol, D) Vehicle control. E) The proportion of memory Th cells expressing IFN-γ is shown as the percentage of IFN-γ^+^ CD4^+^ T cells. F) The proportion of CD4^+^ T cells expressing IL-4 is shown as the percentage of IL-4^+^ CD4^+^ T cells. G) The proportion of CD4^+^ T cells expressing IL-17A is shown as the percentage of IL-17A^+^ CD4^+^ T cells. Expression of H) *TBX21*, I) *GATA3*, and J) *RORC* in RNA extracted from memory Th cells, shown as the relative amounts normalized to housekeeping RNA and compared to the Act group, which was set to 1.0 (dotted line). Pooled data are expressed as Mean ± SEM of five independent biological experiments. One-way ANOVA followed by Tukey’s multiple comparisons tests.

### 3.2. Nebivolol inhibits naïve CD4^+^ T cell shift towards Th17 phenotype when compared to terbutaline

To further illustrate the impact of the β2-AR-biased agonist on the polarization shift of naïve CD4^+^ T cells in the context of Th17 cell development, we initially examined the expression levels of *ADRB1, ADRB2* and *ADRB3* in RNA extracted from naïve Th cells. Naïve T cells expressed *ADRB1* and *ADRB2*, however, *ADRB3* was notably absent (Fig 4A). Subsequently, nebivolol or terbutaline was administered to naïve CD4+ T cells that were activated with Th17-polarizing cytokines and blocking antibodies. Nebivolol demonstrates a significant reduction in IL-17A levels within cell culture supernatants, accompanied by a suppression of RORC expression in Th17-polarized naïve Th cells (Figure 4B, C). Moreover, it effectively decreases the percentage of CD4+ IL-17A+ cells (Figure 4D, E). Conversely, terbutaline exhibits an opposing effect, elevating IL-17A levels, enhancing RORC expression, and increasing the proportion of CD4+ IL-17A+ cells (Fig 4B-E). These findings confirm that β2-AR activation by a biased agonist hinders the shift of human naïve Th cells toward a Th17 phenotype, whereas β2-AR agonist promotes such a shift.

**Fig 4.**
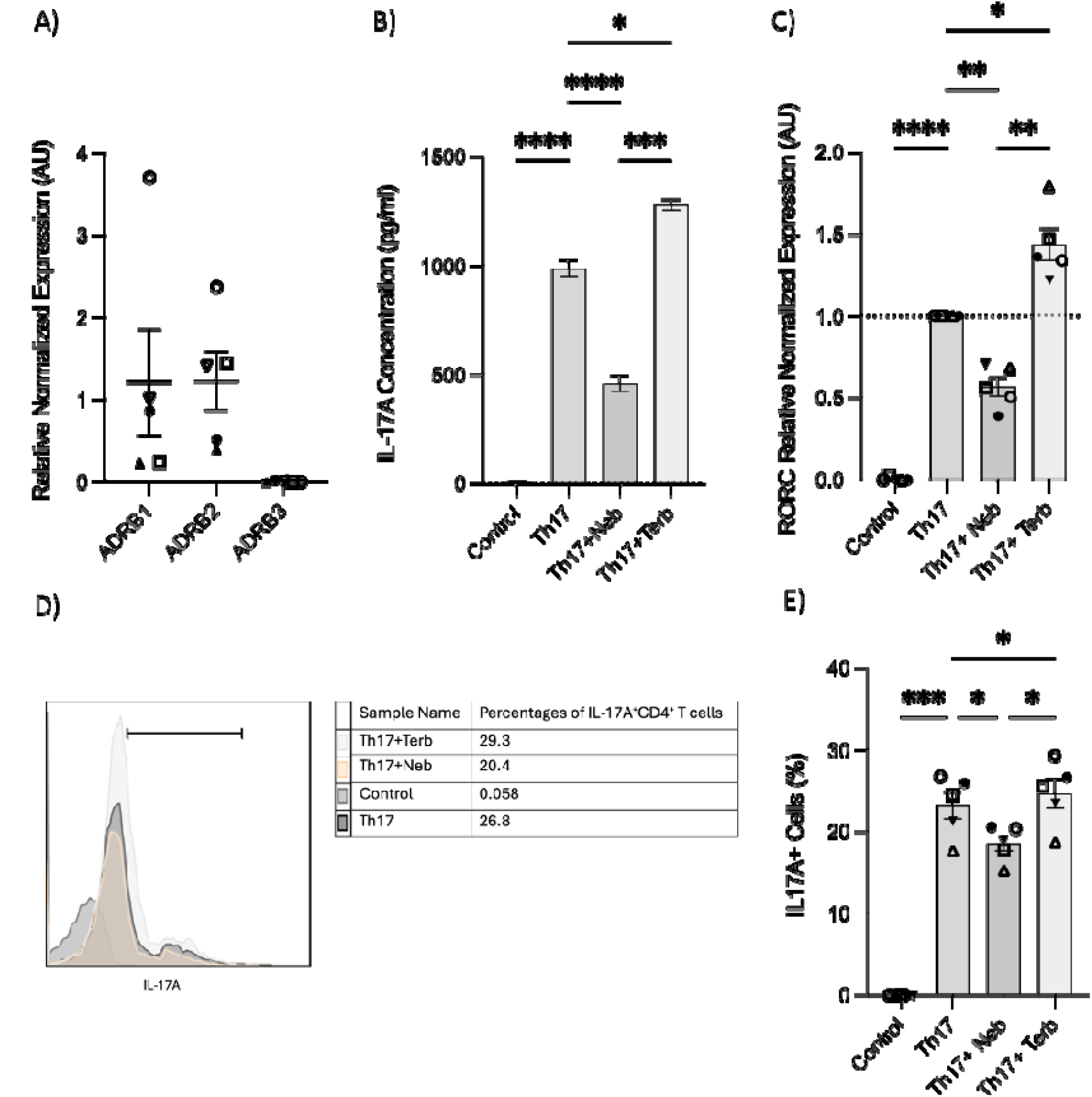
Nebivolol inhibits naïve Th cell shift towards Th17 phenotype, in contrast to terbutaline which augments the shift. A) Expression of *ADRB1*-3 in RNA extracted from naïve CD4^+^ T cells shown as the relative amounts normalized to housekeeping RNA. B) Naïve Th cells were activated with polarizing cytokines and blocking antibodies (Th17) with treatment with nebivolol (Neb) or terbutaline (Terb) for 7 days, and a representative of IL-17A levels in cell culture supernatants were shown. C) RNA expression of the Th17 cell-specific transcriptional factor *RORC* differentiated Th17 cells shown as the relative amounts normalized to housekeeping RNA and compared to shifted Th17 cells, which was set to 1.0 (dotted line). D) A representative of overlapping histogram and E) pooled data of Intracellular cytokine staining shown for CD4^+^IL-17A^+^ cell percentage in polarized Th17 cells after treatment with nebivolol or terbutaline. Pooled data are expressed as the Mean ± SEM of five independent biological experiments. One-way ANOVA followed by Tukey’s multiple comparisons tests.

### 3.3. No significant proliferation and viability effects under nebivolol’s treatment in the memory Th cells

The changes in cytokines and percentages suggested that nebivolol could influence proliferation. However, nebivolol did not change the proliferation index and division index of memory Th cells significantly after treatment (Figure 5A-E). Nebivolol did not change cell viability when compared to activation conditions, as measured by the percentage of necrotic or late apoptotic CD4^+^ T cells with 7AAD after 5 days of cell culture (Figure 5F).

**Fig 5.**
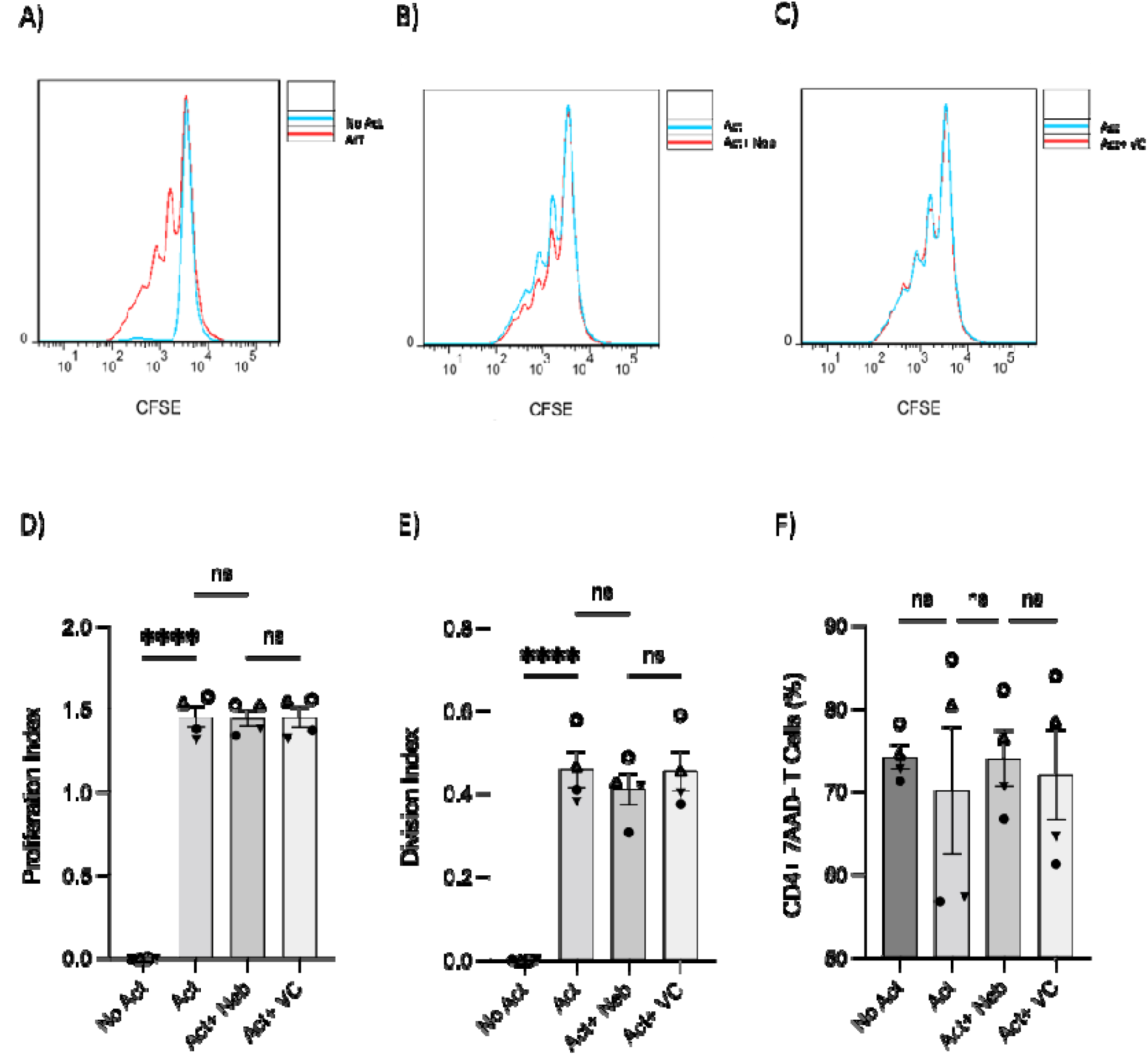
T-cell proliferation and viability were measured by flow cytometry assay. Memory Th cells were not activated or activated with antiCD3/antiCD28/antiCD2 for five days. Representative histograms of the proliferation of memory Th cells in cultures with different groups including A) Activated cells overlayed with non-activated cells, B) Activated cells plus nebivolol overlayed with activation-only condition C) Activated-vehicle control overlayed with activation-only cells. D) The proliferation index of memory Th cells in culture with activation and nebivolol. E) The division index of pooled data. F) The percentage of alive memory CD4^+^, 7AAD^-^ T cells with or without treatment for 5 days gated on all cells in a dot plot by staining with 7AAD in cultures with different groups. Pooled data are expressed as the Mean ± SEM of four independent biological experiments. One-way ANOVA followed by Tukey’s multiple comparisons tests.

### 3.4. Nebivolol’s effect on IL-17A is β2AR-dependent

To investigate receptor involvement, we measured the expression of *ADRB1, ADRB2 and ADRB3* in memory Th cells. Similar to observations in naïve Th cells, the freshly isolated resting memory Th cells expressed *ADRB1* and *ADRB2*, but not *ADRB3 (*Figure 6A). Next, the expression of *ADRB2* in memory Th cells was evaluated after the five-day activation with or without drug treatment. *ADRB2* levels were higher in the non-activated control condition as compared to the activation or activation with nebivolol conditions (Fig 6B). The expression level of β*2AR* protein had a trend to be diminished at the 5-day timepoint which resembles the 5-day mRNA expression data (Fig 6C, D).

**Fig 6.**
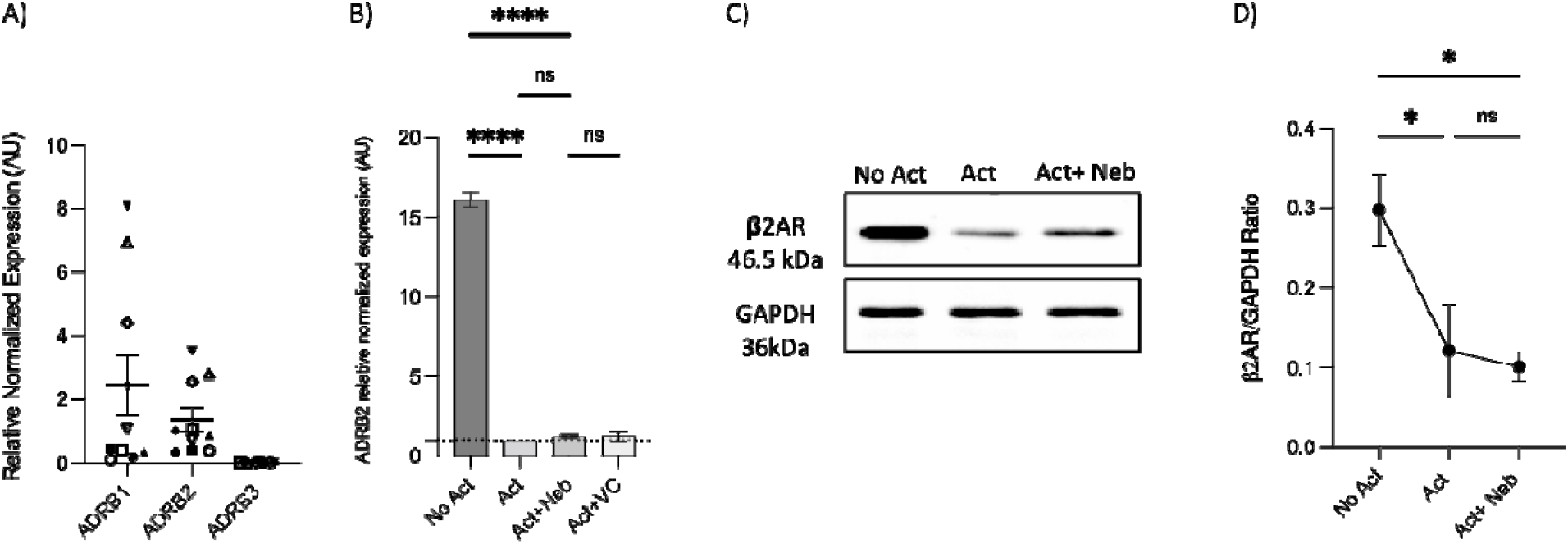
Expression of *ADRB1*-3 in RNA extracted from enrichment memory Th cells and after 5 days with or without treatment. Memory Th cells were not activated or activated with antiCD3/antiCD28/antiCD2 for five days A) Expression of *ADRB1*-3 in RNA extracted from enrichment memory Th cells shown as the relative amounts normalized to housekeeping RNA. Pooled data are expressed as the Mean ± SEM of ten independent biological experiments. B) Expression of *ADRB2* at mRNA levels from Memory Th cells with or without treatment after five days of culture, shown as the relative amounts normalized to housekeeping RNA and compared to the Act group. The data of this figure is representative of three experiments. C) A representative western blot data of equal amounts of protein from the cell lysates was shown for β2AR and GAPDH as a loading control. D) A representative band intensity of five biological experiments was quantified and shown corrected to the loading control as a ratio. The bars show the Mean ± SEM. ANOVA followed by Tukey’s multiple comparisons tests.

Next, we used a pharmacological approach to block receptors in cell culture experiments. The β1AR antagonist (metoprolol) and the pan-β1,2,3AR antagonist (bupranolol) were applied to cell cultures of activated memory Th cells. Metoprolol did not prevent nebivolol from inhibiting IL-17A, in contrast, bupranolol prevented nebivolol from inhibiting IL-17A (Figure 7A, B). To confirm the specific receptor involved in nebivolol’s function, we used electroporation with β2AR-specific multi-sgRNA which significantly reduced β2AR mRNA levels compared to the non-targeting and non-electroporated cells (Control). The viability of cells before and after electroporation was maintained, and the CRISPR/Cas9-mediated knockout of TRAC as a positive control demonstrated over 90% efficiency in mRNA level reduction (Supplementary figures). Quantitative RT-PCR analysis revealed a knockout efficiency of approximately 80%, confirming the successful targeting of the β2AR gene (Figure 7C). Subsequent nebivolol treatment significantly inhibited the β2AR-mediated signaling pathways in β2AR knockout memory Th cells, as indicated by the lack of suppression of IL-17A production compared to the control (Figure 7D). These findings suggest that nebivolol acts in a β2AR-dependent manner, with *ADRB3* not being expressed in the cells, ruling it out as a potential ligand for nebivolol.

**Fig 7.**
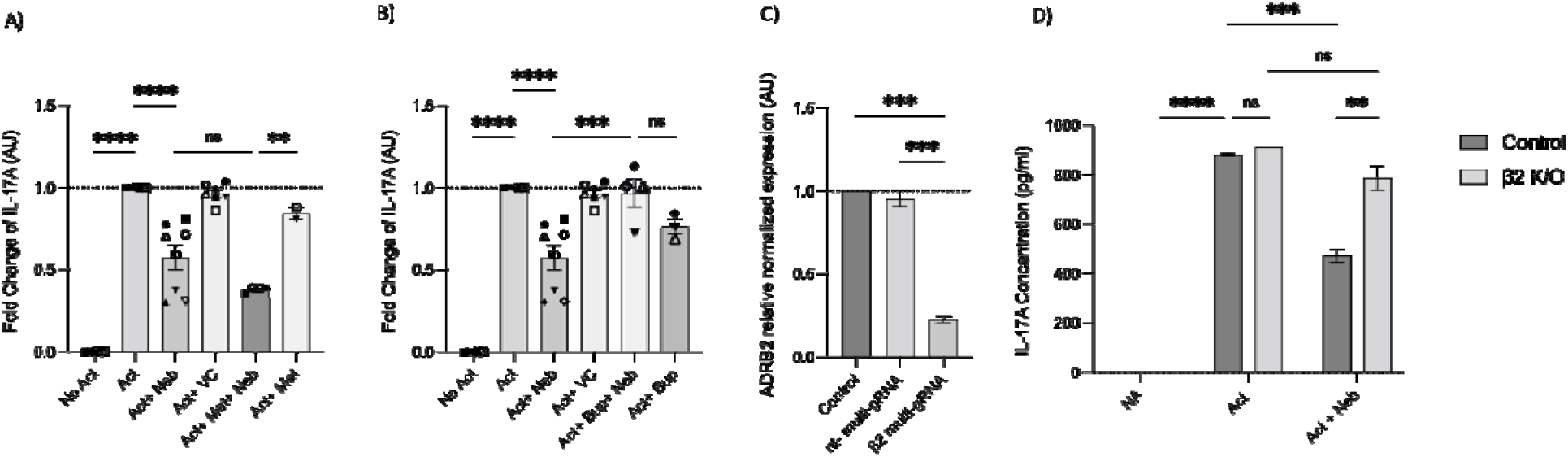
Effect of β-adrenergic receptor antagonists and CRISPR/Cas9-mediated knockout on IL-17A production. Effect of A) β1AR antagonist (metoprolol) and B) non-selective βARs antagonist (bupranolol) on the production of IL-17A on memory Th cells. Activated memory Th cells were blocked for 30 minutes with metoprolol or bupranolol before treatment with nebivolol and incubated for five days. IL-17A was measured in cell culture supernatants. Pooled data are expressed as the Mean ± SEM of four independent biological experiments. Effect of CRISPR/Cas9-mediated knockout of β2AR on nebivolol treatment in activated memory Th cells. Stimulated memory Th cells were electroporated with a CRISPR/Cas9 system targeted by multi-sgRNA against β2AR or non-targeting genes. C) The expression of *ADRB2* at mRNA levels as verification in non-electroporated, non-targeting and β2 multi-sgRNA conditions has shown as the relative amounts normalized to housekeeping RNA and compared to a control set to 1.0 (dotted line). The data of this figure is representative of three independent biological experiments. Three days after electroporation, memory Th cells were cultured with or without nebivolol for another 5 days and D) A representative of IL-17A levels in cell culture supernatants in both control and knock-out β2AR conditions was shown. ANOVA was followed by Tukey’s multiple comparisons tests.

### 3.5. Under non-canonical signaling, nebivolol does not change CREB phosphorylation but promotes the phosphorylation of NF-κB p65

As previously mentioned, our findings indicate that the β2AR agonist terbutaline enhances IL-17A production through a mechanism dependent on PKA and CREB pathways (26). Hypothesizing that nebivolol, acting as a biased agonist, might diverge from this pathway, we investigated its effects on CREB phosphorylation. Memory Th cells were activated and treated with nebivolol or PKA inhibitor H89 plus nebivolol. Nebivolol did not change phosphorylation levels of CREB after 15 minutes, although terbutaline showed a significant increase as was expected (Fig 8A, B). When memory Th cells were co-treated with H89 (a PKA inhibitor) and nebivolol, it did not abrogate the effects of nebivolol on IL-17A (Fig 8C). Thus, the canonical G-protein cAMP-PKA-CREB axis does not appear to be part of nebivolol’s signaling pathway. Given the potential involvement of the NF-κB signaling pathway, we further examined the phosphorylation of NF-κB p65 in activated memory Th cells. Nebivolol inhibited the phosphorylation of NF-κB p65 subunit at position ser529 (Fig 9A, B). Moreover, there was no significant change in phosphorylation of the β-arrestin1-mediated desensitization after 15 minutes of treatment (Supplementary figures). Thus, nebivolol exerts its anti-inflammatory shift by suppressing NF-κB activity possibly through an alternative pathway in memory Th cells.

**Fig 8.**
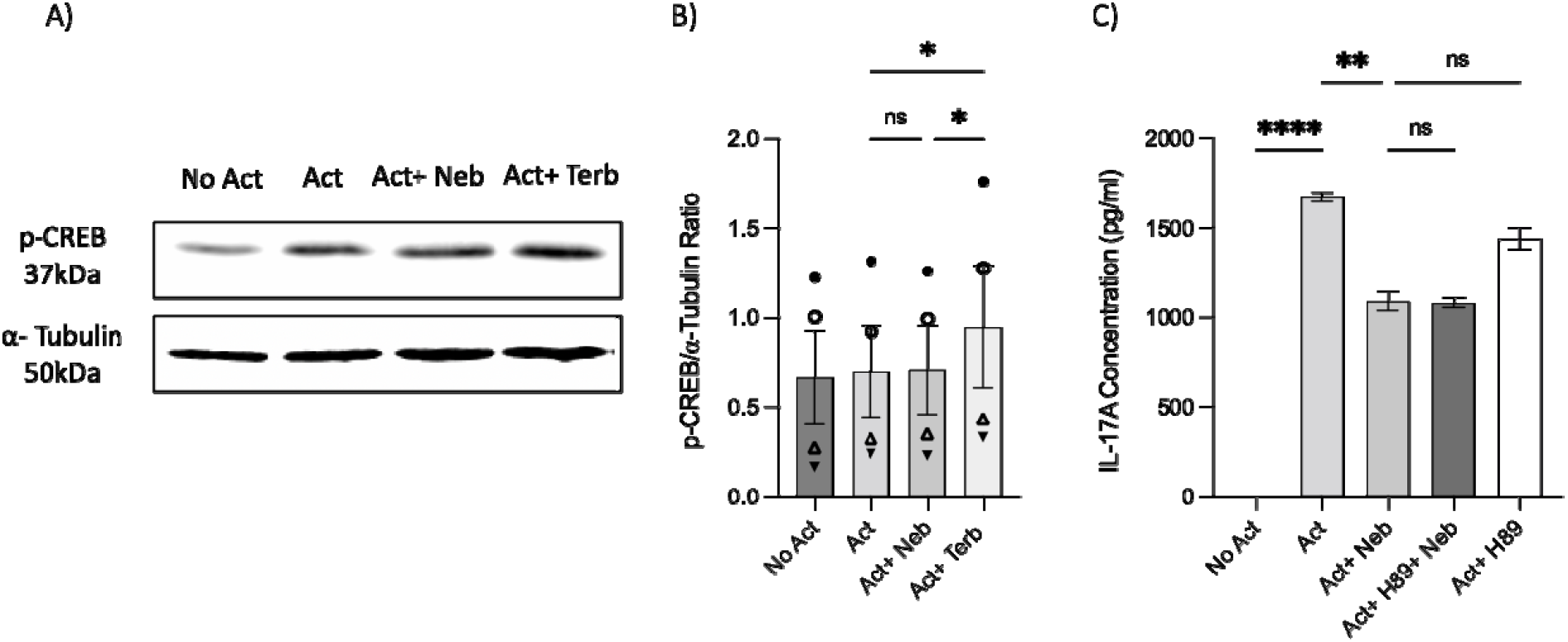
Nebivolol did not stimulate phospho-Ser133-CREB (p-CREB) in memory Th cells. Blasted memory Th cells activated with antiCD3/antiCD28/antiCD2 for 15 minutes in conditions of non-activated cells, activated cells, activated cells plus nebivolol, terbutaline A) Representative western blot data of equal amounts of protein from the cell lysates was shown for p-CREB and α-tubulin as a loading control. B) Band intensity was quantified and shown corrected to the loading control as a ratio, with data pooled from three independent biological experiments. C) Memory Th cells were activated for five days, with nebivolol plus or minus H89. A representative IL-17A ELISA data for the effect of PKA in nebivolol’s signaling pathway by using H89 as a PKA inhibitor. ANOVA was followed by Tukey’s multiple comparisons tests.

**Fig 9.**
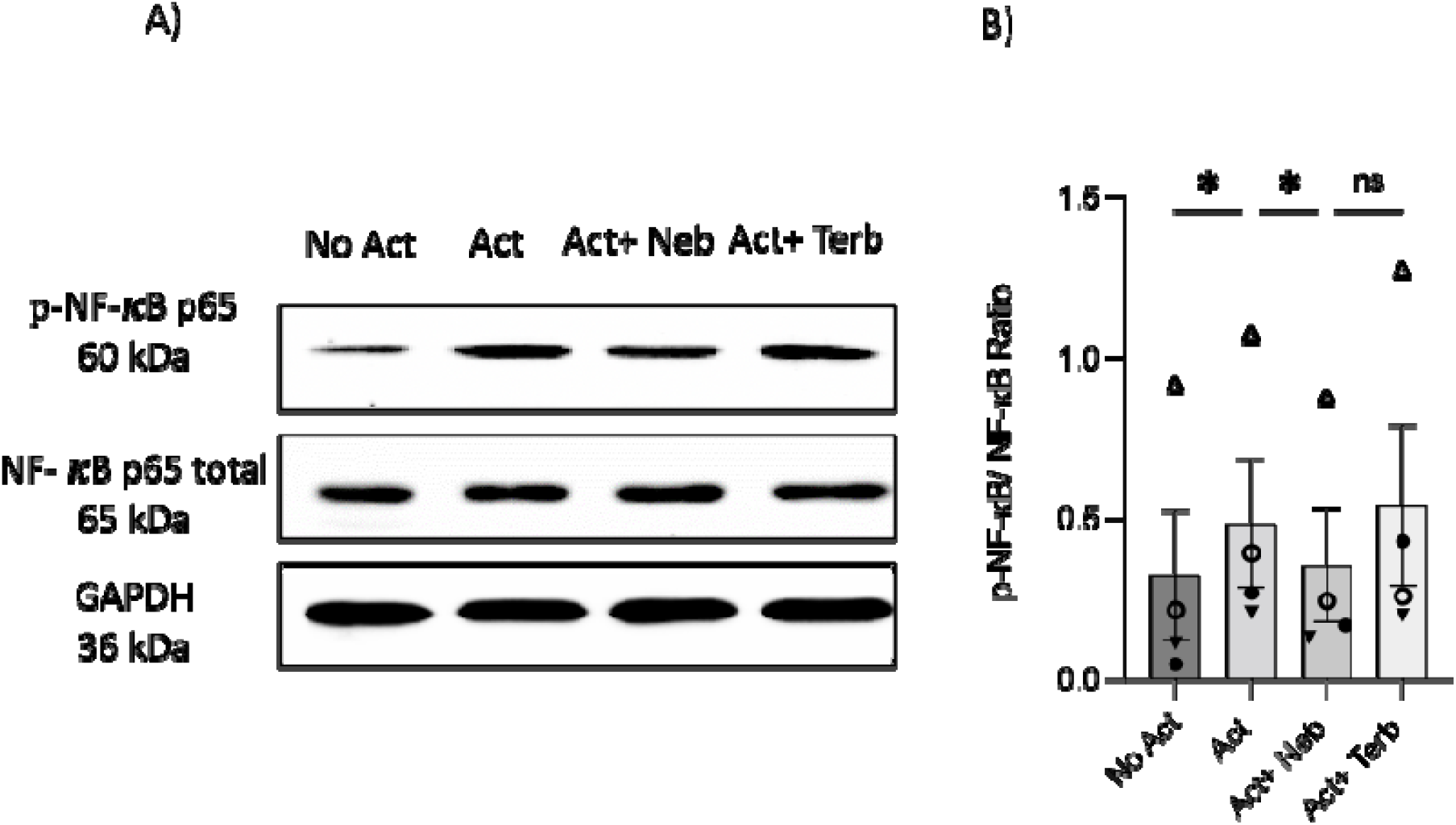
Nebivolol inhibited the phosphorylation of NF-κB p65 in memory Th cells. Blasted memory Th cells were activated with antiCD3/antiCD28/antiCD2 for 15 minutes in conditions of non-activated cells, activated cells, activated cells plus nebivolol. A) Representative western blot data of equal amounts of protein from the cell lysates was shown for p-NF-κB p65 and total NF-κB p65 as a loading control. B) Band intensity was quantified and shown corrected to the loading control as a ratio, with data pooled from four independent biological experiments. ANOVA was followed by Tukey’s multiple comparisons tests.

## Discussion

T cells express adrenergic receptors and respond to adrenergic ligands, but how this impacts pro- or anti-inflammatory cytokine expression is not completely understood. This study demonstrated that nebivolol exerts a notable inhibitory effect on activated naïve and memory Th17 cells, impacting IL-17A secretion levels and the expression of the transcription factor (*RORC*). The data indicates the involvement of a non-canonical NF-κB signaling pathway.

Our initial experiments on PBMCs revealed that nebivolol exhibits notable anti-inflammatory effects by reducing IFN-γ and IL-17A levels while inducing IL-4 production. In contrast, carvedilol did not demonstrate these effects, highlighting the biased and drug-specific nature of nebivolol’s immunomodulatory properties. Since PBMCs are a mixed population, we directed further studies on purified memory and naïve Th cells. Consistent with an anti-inflammatory profile, nebivolol inhibited IFN-γ and augmented IL-4 in the memory Th cells, although the proportions of Th1 and Th2, and the respective transcription factors *TBX21* and *GATA3* did not show significant alterations. This discrepancy between cytokine levels and mRNA expression of transcription factors suggests that potential post-transcriptional or post-translational regulatory mechanisms are at play (38). The drug treatment may influence protein stability, translation efficiency, or cytokine secretion pathways, rather than directly impacting mRNA expression of *TBX21* and *GATA3* which needs further investigation. When examining the levels of IL-10, there was no significant effect of nebivolol. IL-10 is primarily produced by Tregs (28) and has a general anti-inflammatory profile. This lack of effect may stem from a selective impact on certain T cell subsets or signaling pathways. Moreover, we did not assess the proportion of Tregs present in the memory Th cell preparations. Our findings demonstrated that nebivolol mitigated a shift of naïve Th cells toward the Th17 phenotype, whereas terbutaline, acting as a β2-AR agonist, promoted the shift toward Th17 cells. These results imply that nebivolol inhibits Th17 differentiation and function through a distinct mechanism of action, activating β2-AR on CD4^+^ T cells in a manner different from that of a traditional agonist. Overall, nebivolol has the profile of being anti-inflammatory by lowering Th1 and Th17 while augmenting Th2 cell responses.

The study further probed the expression patterns of *ADRB1-3* in isolated naïve and memory human Th cells, shedding light on the receptors’ differential presence in these subsets. Naïve and memory Th cells were devoid of *ADRB3* but expressed *ADRB1* and *ADRB2*. These findings are novel since previous research reported that immune cells express β2-AR predominantly, however memory Th cells were not assessed previously (39). Given nebivolol’s higher affinity for β1-AR as a blocker (30), we introduced the adrenergic receptor antagonists into the experimental setup to determine which receptor was mediating the effect. The distinct responses observed with the β1AR antagonist and non-selective βAR antagonist emphasize the unique anti-inflammatory effects of nebivolol, consistent with previous findings (40). Using a CRISPR/cas9 method of genome targeting *ADRB2* which enables rapid and flexible experimental manipulation (41), we achieved an 80% knockout efficiency in human memory Th cells. This demonstrated that nebivolol’s inhibitory function on IL-17A production is mediated through the β2AR pathway.

Our investigation into the expression of β2AR following activation and treatment over 5 days, at both mRNA and protein levels, revealed a notable reduction in receptor expression in memory Th cells treated with or without nebivolol compared to the non-activation group. This decrease in β2AR expression may be attributed to receptor desensitization induced by stimulation of the T cell receptor (TCR) and β2AR signaling pathways. This feedback loop involves several steps leading to a decrease in receptor quantity on both the cell surface and intracellular compartments. These steps include the degradation of internalized receptors and the promotion of mRNA decay (42, 43), indicating the involvement of transcriptional control mechanisms (44).

The significant inhibition of NF-κB p65 activation by nebivolol, while CREB phosphorylation remained unaffected, suggests a divergence from the canonical β2-adrenergic receptor (β2AR) signaling pathway. In canonical signaling, activation of protein kinase A (PKA) typically facilitates CREB phosphorylation, competing with NF-κB activation and impeding its transcriptional activity (45). However, PKA activation of cytoplasmic kinases, such as p38 mitogen-activated protein kinase (MAPK), enhances IκBα degradation, thereby activating NF-κB-driven transcription (46). Notably, our results indicate that the β2AR agonist drug terbutaline, unlike nebivolol, facilitated CREB phosphorylation along with NF-κB activation, consistent with prior research in other types of T cells (47, 48). These nuanced differences underscore the complexity of β2AR-mediated pathways and suggest nebivolol may act through a non-canonical pathway.

While we measured β-arrestin-1, our study did not measure β-arrestin 2 activation or its potential involvement in nebivolol’s effects. β-arrestin 2 is known to inhibit NF-κB activation and the expression of its target genes by binding to and inhibiting the degradation of IκBα (14). Our findings provide insight into nebivolol’s mechanism of action, suggesting a β-arrestin biased agonism that selectively modulates downstream signaling molecules. This may explain nebivolol’s suppressive impact on IL-17A, independent of alterations in T-cell proliferation or viability. Thus, nebivolol appears to modulate cell signaling pathways relevant to cytokine expression without affecting the cell cycle or causing cellular cytotoxicity.

The observed outcomes align with established knowledge about adrenergic signaling in immune cells, emphasizing the role of AR in regulating T-cell function. β2-ARs, expressed widely throughout the body, including immune and non-immune cells, play a bystander role in immune responses, making them an optimal focal point for studying immune control. While β2-AR agonists have been recognized as modulators of inflammatory responses, β2-AR biased agonists (49), like nebivolol, selectively stimulate specific pathways, offering a more targeted approach to modulating immune cell function and treating disease (50). Nebivolol, identified as a notable β2-biased agonist in this study, emerges as a promising therapeutic target for chronic inflammation. Recent findings indicate its efficacy in reducing inflammatory signs of psoriatic lesions, inhibiting serum inflammatory biomarkers like IL-17A and TNF-α which was independent of β1-blockade activity (51, 52). The significance of nebivolol lies in its antioxidant and anti-inflammatory potential, providing a promising avenue for addressing persistent health concerns through the modulation of immune responses and cytokine production.

The immunomodulatory properties of nebivolol, particularly its ability to inhibit pro-inflammatory Th17 cell responses and promote anti-inflammatory Th2 responses, suggest promising therapeutic applications in autoimmune and inflammatory diseases such as multiple sclerosis, rheumatoid arthritis, and psoriasis characterized by elevated Th17 response (53, 54). Moreover, memory Th cells hold significant relevance in autoimmune diseases due to their enduring nature and ability to orchestrate recurrent autoimmune reactions (18). Memory Th17 and Th1 cells in the blood of MS patients correlate with disease severity (55). The established safety profile of nebivolol as a cardiovascular drug enhances its potential for repurposing, offering a well-tolerated option that could complement existing therapies. Future research should focus on elucidating the mechanisms underlying nebivolol’s biased agonism and assessing its long-term efficacy and safety in chronic inflammation, which could pave the way for innovative treatment strategies in immune modulation.

Our study acknowledges several limitations. The small sample size may limit the generalizability of our findings, necessitating larger cohorts in future research. Assessing specific molecules within the β-arrestin signaling pathway could confirm nebivolol’s biased agonist properties. Additionally, evaluating nebivolol’s effects in vivo in inflammatory situations would provide valuable insights into its therapeutic potential. The lack of transcriptomic analyses such as RNA sequencing is a limitation that could be addressed in future studies to explore the broader molecular effects of nebivolol. Addressing these limitations will build on our findings and deepen our understanding of nebivolol’s role in immune modulation.

In conclusion, this study advances our understanding of nebivolol’s immunomodulatory potential by specifically elucidating its impact on IL-17A secretion in Th17 cells and the underlying mechanisms. Nebivolol emerges as a promising candidate for further exploration in immunopharmacology, leveraging its established safety profile through drug repurposing. These findings offer avenues for targeted therapeutic interventions in chronic inflammation management, potentially reshaping treatment approaches with a familiar and accessible tool.

## Acknowledgements

This work was funded by NSERC Discovery grant RGPIN□2019□ 06980 and FRNQT doctoral fellowship in the research grant 2023-2024 - B2X – 334802.

