## Supplemental Figures for "β2-Adrenergic Biased Agonist Nebivolol Inhibits the Development of Th17 and the Response of Memory Th17 Cells in an NF-κB-Dependent Manner"

### 1 **Supplementary Data**

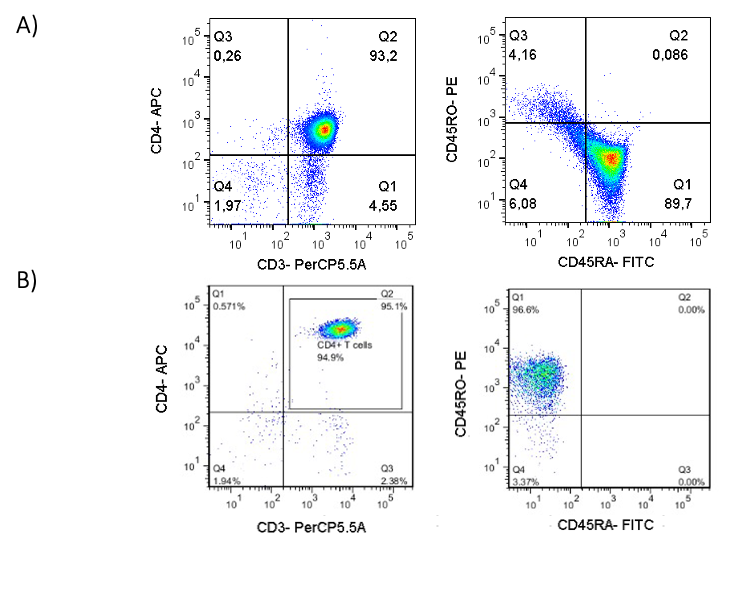

**Supplementary Figure 1.** Purification checks of A) naïve and B) memory Th cells after using EasySep® naïve and memory CD4+ T cell enrichment kits respectively.

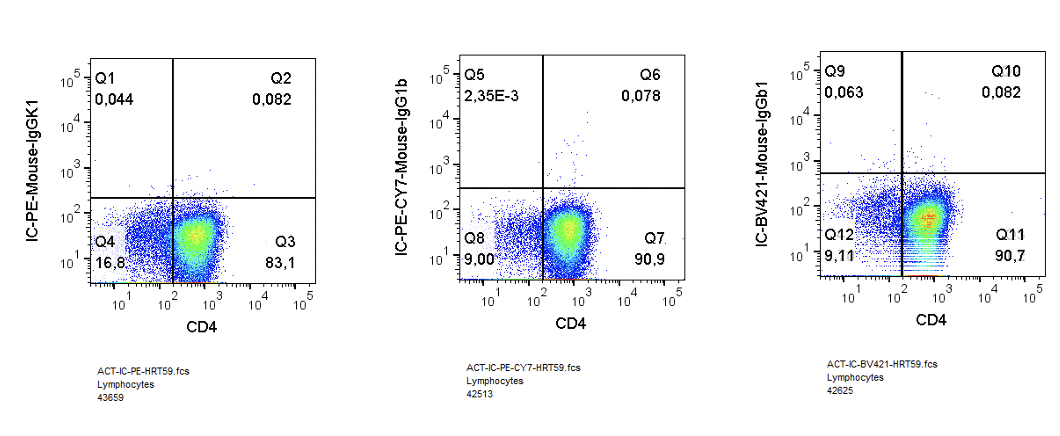
**Supplementary Figure 2.** Isotype controls of PE-Mouse-IgGk1 (IL-17A), PE-Cy7-Mouse-IgG1b (IL-4), and BV421-Mouse-IgG1b (IFN-γ) correspond to the type and isotype of mAbs used in intracellular cytokine staining (ICS). The same gating strategy was applied for each experiment presented in the manuscript.

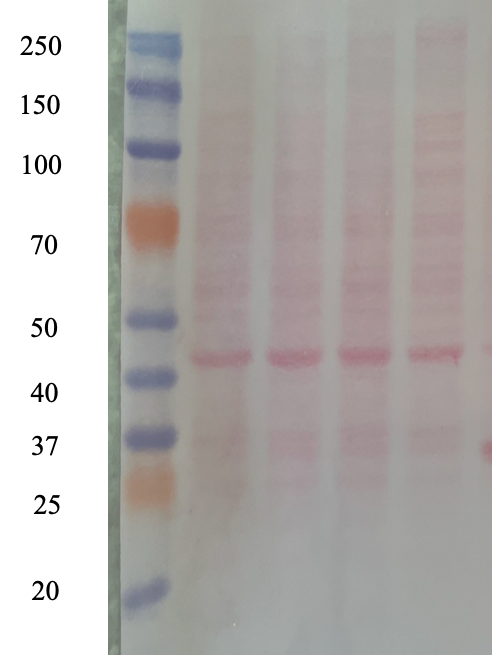

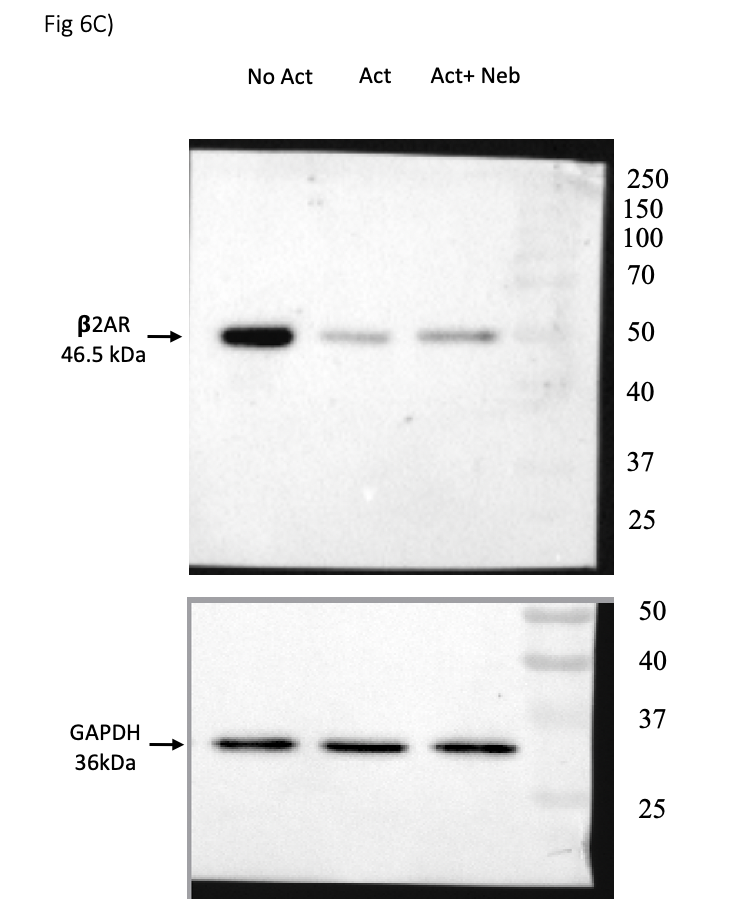

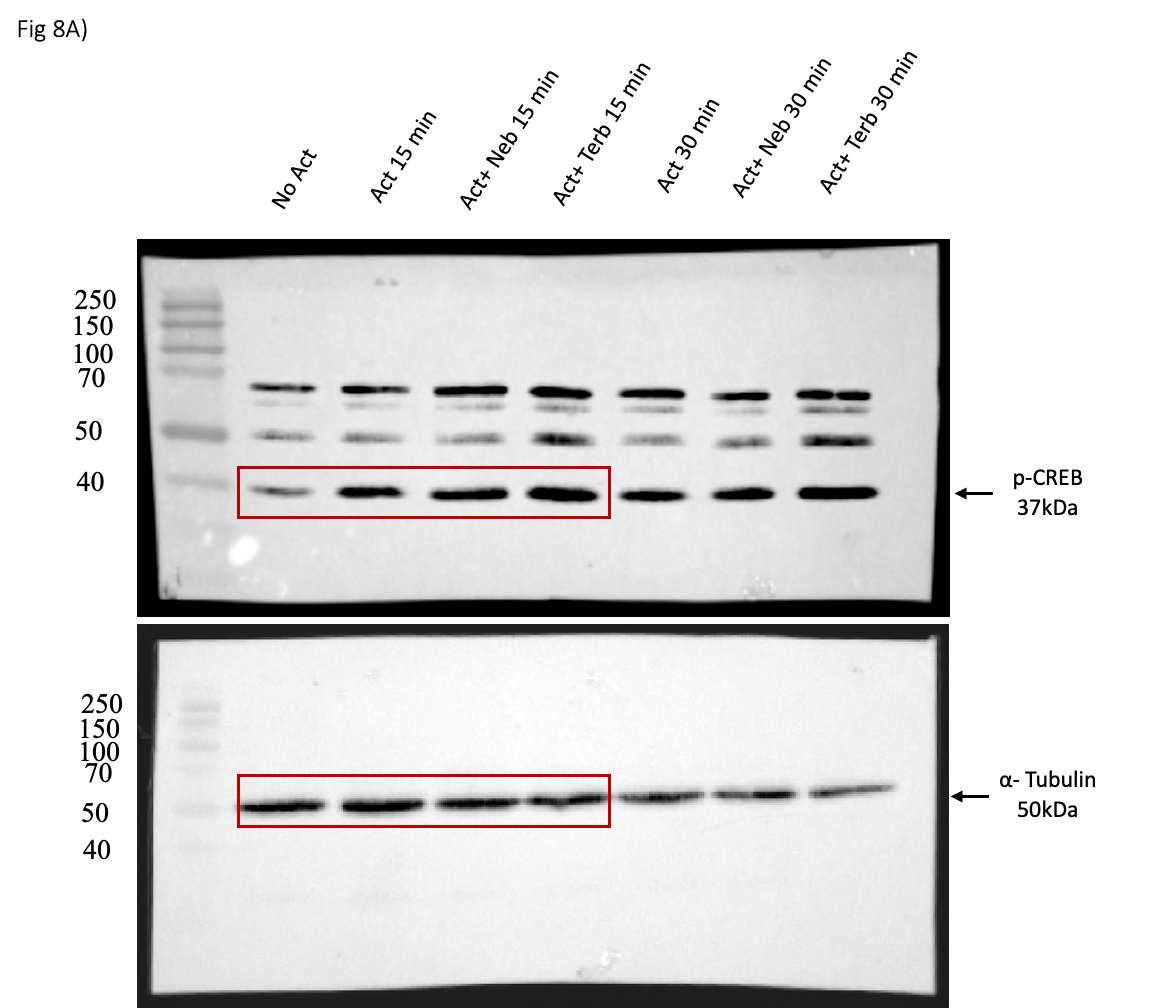

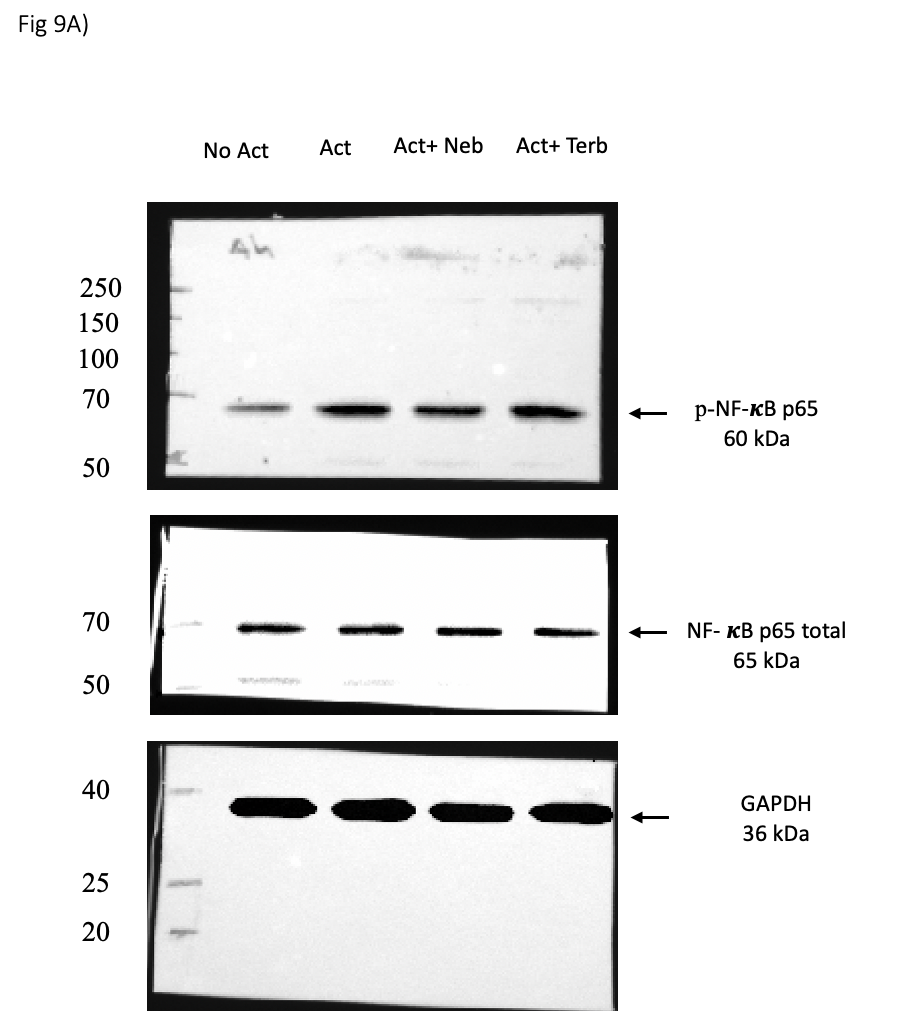

**Supplementary Figure 3.** Images of unprocessed nitrocellulose blotting membranes, including the Precision Plus Protein Dual Color Standards ladder, are presented alongside the full western blots for Figures 6C, 8A, and 9A, respectively. These images provide the uncropped western blot data, ensuring transparency and confirming the consistency of the bands observed in the manuscript figures. This allows for the verification of protein sizes and loading controls across the experimental conditions.

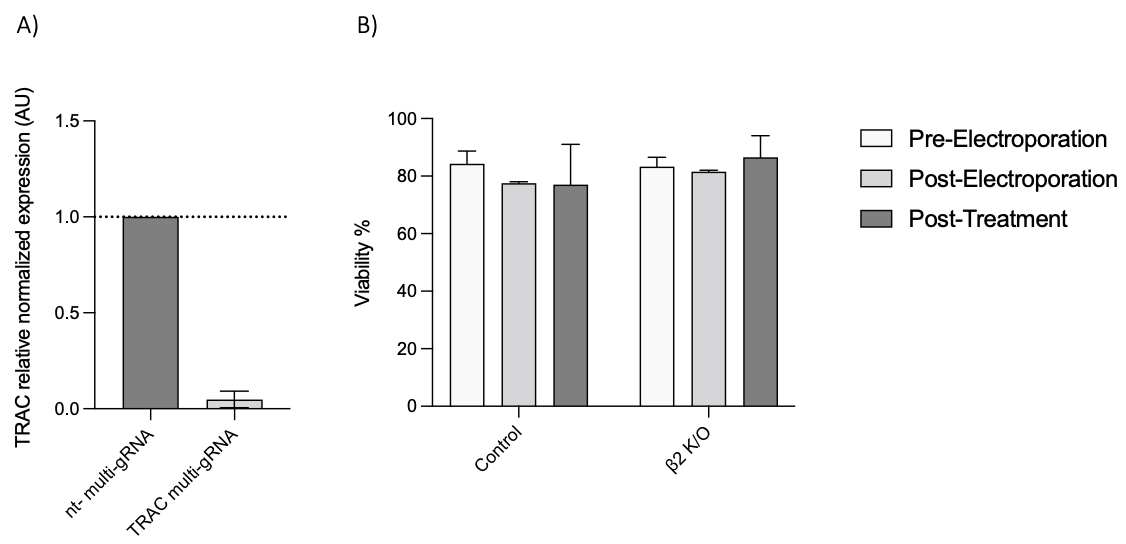

**Supplementary Figure 3.** A) Verification of CRISPR/Cas9 knockouts of the TRAC gene as a positive control in memory Th cells at the mRNA level. The graph includes non-targeting (nt) and TRAC multi-sgRNA conditions. Results are shown as relative amounts normalized to housekeeping RNA, compared to nt-multi-gRNA, set to 1.0 (dotted line). B) Viability assessment of non-electroporated and ADRB2 multi-sgRNA conditions before and after electroporation on DAY 4 and after 5 days of treatment. Data represent the mean ± SEM from two experiments. Statistical analysis was performed using one-way ANOVA followed by Tukey’s multiple comparisons test.

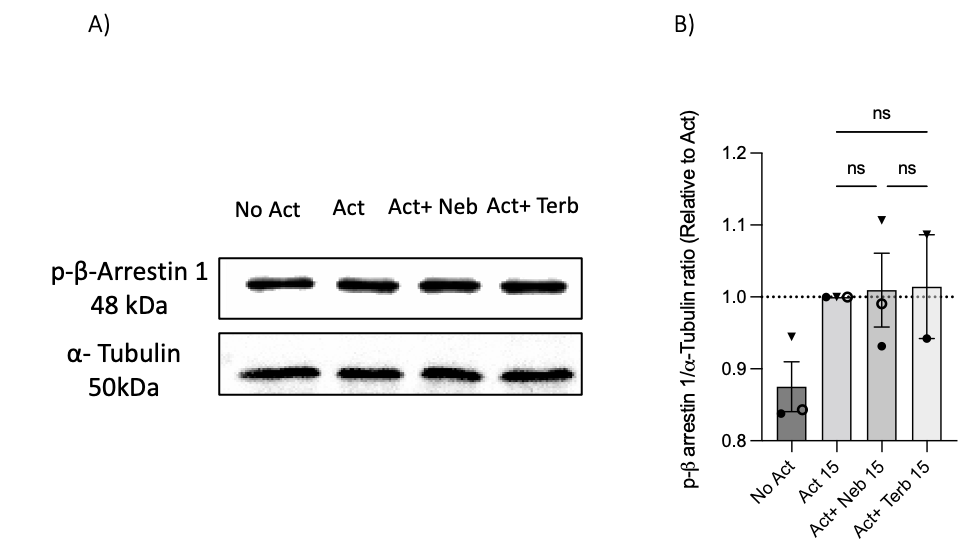

**Supplementary Figure 4.** Nebivolol did not stimulate phospho-beta-arrestin-1 (p-ß Arrestin1) in memory Th cells. Blasted memory Th cells were activated with ImmunoCult for 15 mins in conditions of non-activated cells, activated cells, activated cells plus nebivolol, or H89 plus nebivolol A) Representative western blot data of equal amounts of protein from the cell lysates was shown for p-ß arrestin1 and α-tubulin as a loading control. Band intensity was quantified and shown corrected to the loading control, and relative to the non-activation, which was set to 1 for p-ß arrestin1 (dotted line), with data pooled from 3 experiments.

**Supplementary Table 1,** payload sequence of multi-guide RNA for targeting TRAC and ADRB2 genes.

| Human TRAC multi-guide sgRNA, modified | sgRNA 1: 5'-CUCUCAGCUGGUACACGGCA-3' |
| --- | --- |
|  | sgRNA 2: 5'-GAGAAUCAAAAUCGGUGAAU-3' s |
|  | gRNA 3: 5'-ACAAAACUGUGCUAGACAUG-3' |
| Human ADRB2 multi-guide sgRNA, modified | sgRNA 1: 5'-GCCGUUCCCGGGUUGCCCCA-3' |
|  | sgRNA 2: 5'-ACCACGACGUCACGCAGGAA -3' |
|  | gRNA 3: 5'-CGUCAUGUCUCUCAUCGUCC -3' |
